# Travelling spindles create necessary conditions for spike-timing-dependent plasticity in humans

**DOI:** 10.1101/2020.05.21.079152

**Authors:** Charles W. Dickey, Anna Sargsyan, Joseph R. Madsen, Emad N. Eskandar, Sydney S. Cash, Eric Halgren

## Abstract

Sleep spindles facilitate memory consolidation in the cortex during mammalian non-rapid eye movement (NREM) sleep. In rodents, phase-locked firing during spindles may facilitate spike-timing-dependent plasticity (STDP) by grouping pre- and post-synaptic cell firing within ∼25ms. Currently, microphysiological evidence in humans for conditions conducive for STDP during spindles is absent. We analyzed local field potentials and supragranular unit spiking during spindles from 10×10 arrays of microelectrodes at 400µm pitch in humans. We found strong tonic and phase-locked increases in firing and co-firing within 25ms during spindles. Co-firing, spindle co-occurrence, and spindle coherence were greatest between sites within ∼2mm, and high co-firing of units on different electrodes was largely restricted to moments of high spindle coherence between those electrodes. Spindles propagated at ∼0.23m/s in distinct patterns, with correlated cell co-firing sequences. These results suggest that spindles may organize spatiotemporal patterns of neuronal co-firing which promote memory consolidation during NREM sleep.

## Introduction

Sleep spindles are bursts of 10-16Hz oscillations that last for 0.5-2s at the scalp and occur spontaneously during mammalian non-rapid eye movement (NREM) sleep. Spindles are generated by an interaction of intrinsic currents and local circuits within the thalamus (Destexhe and Sejnowski, 2003; McCormick et al., 2015), and are projected to all cortical areas (Piantoni et al., 2017; Mak-McCully et al., 2017). Spindles often occur on up-states following down-states (Gonzalez et al 2018; Mak-McCully et al., 2017).

In the two-stage model of memory, short-term memories are encoded in the hippocampus and then subsequently consolidated into long-term storage by repeated activation of cortical networks during sleep (Buzsáki et al., 1998; McClelland et al 1995). It is hypothesized that hippocampal-to-cortical transfer of memories involves the replay of cell-firing sequences during sleep (Skaggs and McNaughton, 1996), correlated with hippocampal sharp-wave ripples, cortical slow oscillations, and cortical sleep spindles (Jiang et al., 2019a; Jiang et al., 2019b, Jiang et al., 2017; Buzsáki, 2015; Johnson et al., 2010). Disrupting this association in rodents impairs consolidation (Maingret et al., 2016), and spindle density is correlated with consolidation in humans (Mednick et al., 2013; Cox et al., 2012), suggesting that cortical spindles contribute to the consolidation process (Diekelmann and Born, 2010). However, the mechanisms underlying this phenomenon are not well understood.

In rodents, spindles are associated with dendritic Ca2+ influxes (Seibt et al., 2017), which could promote plasticity during memory consolidation. In support of this hypothesis, *in vitro* stimulation of rat cortical pyramidal cells pattern matched to *in vivo* firing sequences during spindles promotes Ca2+-dependent long term potentiation (LTP) of excitatory postsynaptic potentials (Rosanova and Ulrich, 2005). Potentiation was dependent on coordinated pre-synaptic potentials and post-synaptic spiking. These observations suggest that spindles may promote spike-timing-dependent plasticity (STDP), which is a Ca2+-dependent mechanism where correlated pre- and post-synaptic spiking within a short time window modulates synaptic strength (Feldman, 2012). In the standard model, STDP facilitates LTP when pre-synaptic spiking occurs within 25ms before post-synaptic spiking, and long term depression (LTD) when the post-synaptic cell fires first.

Indirect evidence in humans supports the hypothesis that spindles facilitate STDP. Specifically, electrocorticography recordings show that spindles travel across the cortex at ∼3-9m/s, which is optimal for inducing STDP across distant regions (Muller et al., 2016). Furthermore, intracranial studies show that spindles are associated with increased high gamma, which may reflect increased cortical unit spiking (Hagler et al., 2018; Gonzalez et al., 2018) that could lead to increased co-firing, required for STDP. However this remains controversial as a previous human intracranial study did not find an increase in unit spiking during spindles (Andrillon et al. 2011). While several limitations preclude the recording of dendritic Ca2+ currents in humans, it is possible to test if the unit spike timing requirements for STDP are fulfilled during spindles. Specifically, the most prominent requirement is that STDP requires co-firing within 25ms.

We analyzed intracranial microelectrode recordings from cortical supragranular layers II/III, and possibly as low as layer IV, in patients undergoing evaluation of pharmaco-resistant intractable epilepsy. We detected unit spikes on each of the 96 channels in the microarray and classified units as putative pyramidal (PY), interneuron (IN), or multi-unit (MU) based on standard neurophysiological criteria (Barthó et al., 2004; McCormick et al., 1985). We detected spindles on these same channels in the local field potential (LFP). Spindles were associated with both tonic and phase-specific increases in unit firing as well as ordered co-firing within 25ms, which fulfils a critical precondition for STDP. Some unit pairs had a preferred order of firing, which could support directional plasticity. Spindles tended to co-occur and were highly coherent within ∼1.5-2.0 mm, and increased co-firing of units on different electrodes was highly enriched when those electrodes displayed highly-coherent spindles. Spindles, and associated co-firing, propagated at ∼0.23ms with multiple patterns on a given array and within the same spindle. Thus, spindles spatiotemporally organize neuronal co-firing on a sub-centimeter scale in a manner that could facilitate plasticity across multiple networks.

## Methods

### Participants and Data Collection

Four adult patients (**Table 1**) with focal, pharmaco-resistant epilepsy underwent 4-21 days of continuous electrocorticography and invasive EEG recordings for the localization of seizure foci prior to resection. The decision to implant and the duration of implantation were based entirely on clinical grounds. While undergoing clinical recording these patients also underwent intracranial microelectrode recordings with the Utah Array (**Fig.1A;** Utah Array – © 2020 Blackrock Microsystems, LLC). In all cases the Utah Array was implanted in a location that was strongly suspected based on pre-implant clinical information to be included within the boundary of the therapeutic resection, and in all cases it was later resected (**Fig.1B**). This was done for research purposes and did not disrupt clinical monitoring. Patients agreed to participate in these research studies after fully informed consent according to the Declaration of Helsinki guidelines as monitored by the local Institutional Review Boards.

**Figure 1:**
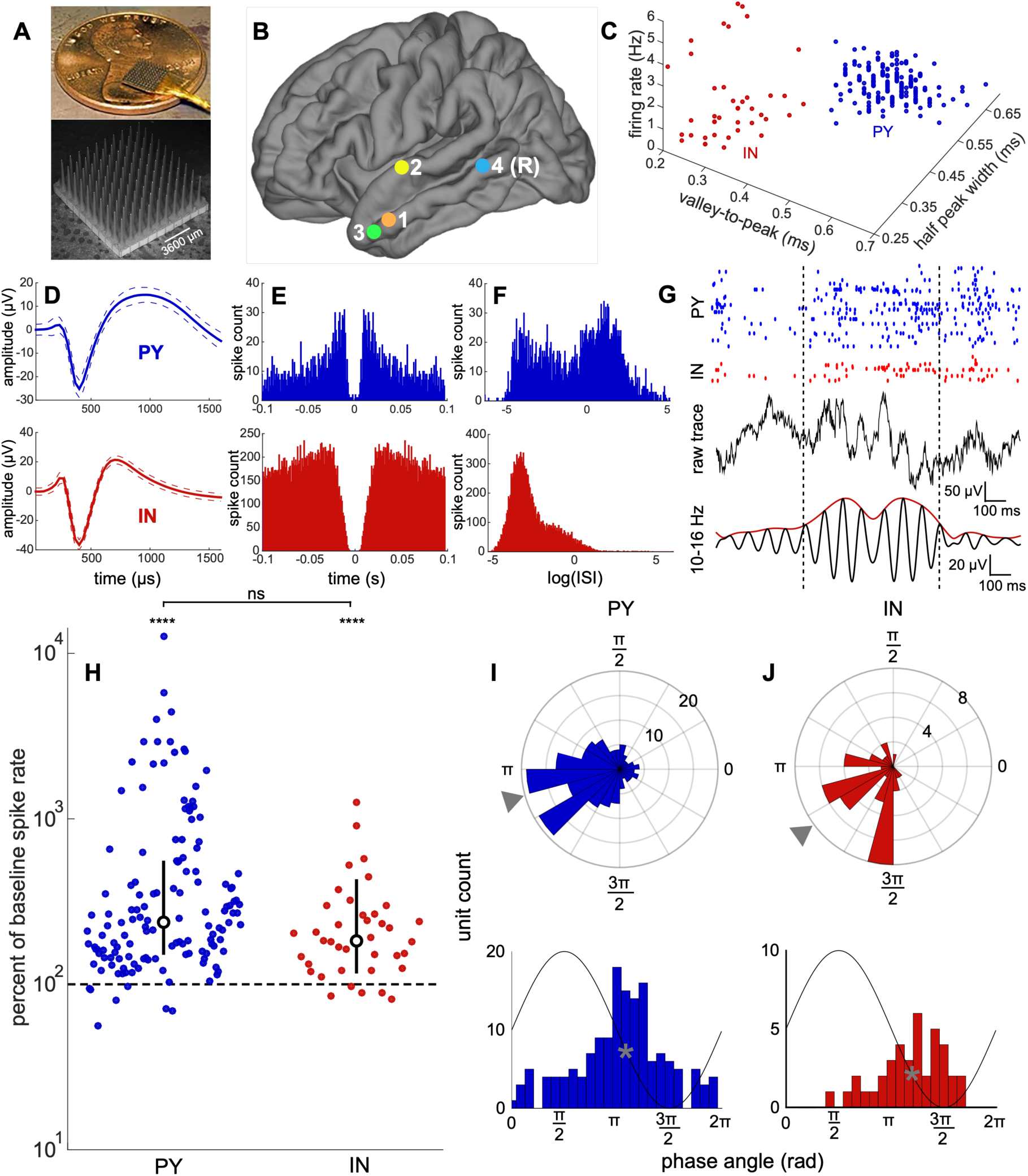
Array implantation, single unit classification, spindle detection, and unit spiking during spindles. **A**, images of the Utah Array. **B**, Utah Array implantation locations for all 4 patients. All sites of implantation were on the left except for patient 4, which was on the right (R). **C**, PYs (blue) and INs (red) form separate clusters based on waveform half-width duration, waveform valley-to-peak duration, and firing rate. **D-F**, average and standard deviation spike waveform (**D**), spike autocorrelation (**E**), and ISI distribution (**F**) for an example PY and IN. **G**, raw and 10-16Hz bandpassed traces of a sleep spindle with raster plot of associated unit spiking of the three putative cell types. **H**, firing rates during spindles for PYs and INs as log-percent of baseline firing rates. The spindle spike rate of each unit was computed as the firing rate across all spindles. The baseline spike rate was computed as the spike rate during NREM intervals between spindles. Each colored circle shows the mean spike rate of one unit. White circle shows the median value and error bars show the 25th and 75th percentiles. Dashed horizontal line shows 100% percent, which is equal to the baseline spike rate. **I-J**, polar and non-polar histograms show circular mean spindle phases of spiking of PYs (**I**) and INs (**J**). One cycle of a spindle is superimposed on non-polar histograms to visualize the phase-spike timing relationship. Gray triangles on polar histograms and gray asterisks on non-polar histograms show circular means. Bonferroni-corrected ****p<0.0001. IN=putative interneuron unit, PY=putative pyramidal unit, ISI=inter-spike interval.

**Table 1:** Patient demographics, array implantation locations, and unit characteristics. The total units and total spikes for each unit type across all four patients are reported. The average and standard deviation across units for all patients for peak-to-valley, firing rate, peak-to-valley width, half peak width, and bursting index are reported. PY=pyramidal unit, IN=interneuron unit, MU=multi-unit.

| Patient Demographics and Array Implantation |  |  |  |  |  |  |  |
| --- | --- | --- | --- | --- | --- | --- | --- |
| Patient | Age | Sex | Handedness | Utah Array implantation location |  | Probe length (mm) |  |
| 1 | 51 | F | R | Left middle temporal gyrus |  | 1.0 |  |
| 2 | 31 | M | L | Left superior temporal gyrus |  | 1.5 |  |
| 3 | 47 | M | R | Right middle temporal gyrus |  | 1.5 |  |
| 4 | 21 | M | R | Left middle temporal gyrus |  | 1.0 |  |
| Unit Characteristics |  |  |  |  |  |  |  |
|  | Total Units | Total Spikes | Peak-to-Valley Amplitude (μV) | Firing Rate (Hz) | Peak-to-Valley Width (ms) | Half Peak Width (ms) | Bursting Index |
| PY | 156 | 352992 | 84.02 ± 59.45 | 0.19 ± 0.17 | 0.49 ± 0.057 | 0.61 ± 0.038 | 0.046 ± 0.033 |
| IN | 39 | 785571 | 41.60 ± 26.09 | 1.71 ± 1.70 | 0.30 ± 0.054 | 0.35 ± 0.052 | 0.012 ± 0.016 |
| MU | 158 | 5256429 | 26.53 ± 14.89 | 2.82 ± 2.68 | 0.48 ± 0.086 | 0.57 ± 0.070 | 0.046 ±0.033 |

### Electrodes and Localization

The Utah Array is a 10×10 microelectrode grid, with corners omitted, that has 400μm contact pitch (**Fig.1A**). Each silicon probe is 1 or 1.5mm long (summarized in **Table 1**) and 35-75μm wide at its base, tapering to 3-5μm at the tip, and is insulated except for the tip, which is platinum-coated. For our studies the array was implanted into the superior or middle temporal gyrus, and not within the bounds of the epileptogenic focus, by the neurosurgeon under direct visualization such that the probes were perpendicular to the cortical surface. Based on a previous histological examination of human brain tissue, temporal cortex layer II begins at an average of 252µm and layer III ends at an average of 1201µm (Mohan et al., 2015). Therefore, we expect that the 1.0-1.5mm long electrodes of the Utah array were implanted in supragranular layers II/III and possibly as low as upper granular layer IV.

### Recording and Preprocessing

Data were acquired at 30kHz sampling (Blackrock Microsystems), from 0.3 to 7.5kHz. Data were subsequently low-passed at 500Hz and down-sampled to 1kHz for the LFPs. Data were saved for offline analysis using custom-written scripts in MATLAB 2019b (MathWorks). LFPs were visualized in MATLAB: *FieldTrip* (Oostenveld et al., 2011). Channels were excluded when there were large amounts of noise or no units detected. Out of the 96 recording channels the average number excluded from analysis was 29.75 (range 13-47). The 1kHz data was average-referenced to negate the effects of the distant subdural reference, which could have detected neural activity distant from the array.

### Sleep Staging

After the data were collected, wake, NREM sleep, and REM sleep periods were determined based on visual examination of electrocorticography data by clinicians with expertise in sleep staging. Periods marked as NREM stages 2 and 3, based on the presence of slow waves, K-complexes, and spindles, were selected for analysis. These periods were validated as NREM stages 2 and 3 based on increases in delta and sigma band powers.

### Spike Detection and Sorting

The 30kHz data recorded from each electrode contact was bandpassed at 300-3000Hz with an 8th order elliptic filter with a pass-band ripple of 0.1dB and a stop-band attenuation of 40dB. Putative unit spikes were detected when the filtered signal exceeded 5 times the estimated standard deviation of the background noise (Donoho and Johnstone, 1992), computed as 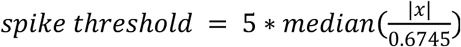 where *x* is the 300-3000Hz bandpassed data. The first three principal components of each spike were computed and unit clusters were manually selected. The remaining data points underwent clustering by k-means and a Kalman filter mixture model (Calabrese and Paninski, 2011). Visual inspection of the clusters identified by these two algorithms was used to determine which achieved better separation. Spikes were examined visually and those with abnormal waveform shapes or amplitudes far exceeding the majority of the spikes from their putative unit, such as those that may have been due to epileptiform activity, were excluded from analysis.

### Unit Classification

PYs fire at low rates (∼0.1Hz), with frequent bursting, and have short refractory periods and sharp spike autocorrelations, whereas INs typically fire at high rates (>1Hz), with infrequent bursting, and have long refractory periods and broad spike autocorrelations. We classified units based on established methods in rodents (Barthó et al., 2004; McCormick et al., 1985) that have been adapted for use in humans (Peyrache et al., 2012), with additional considerations for MUs, which had spikes that exceeded the detection threshold but could not be clustered into separable units. For each unit we computed the firing rate, valley-to-peak time interval, half peak width time interval, and bursting index (summarized in **Table 1**; **Fig.1C** shows independent clusters of PY and IN based on firing rate, valley-to-peak interval, and half width interval). As bursting results in a bimodal distribution of inter-spike intervals (ISIs), the bursting index was determined by running the Hartigan dip test for unimodality on the logarithm of distribution of ISIs (Hartigan et al., 1985). Units were classified as putative PYs if they had spike rates of ∼0.1-0.8Hz, long valley-to-peak and half width intervals (**Fig.1D**), sharp autocorrelations (**Fig.1E**), and a bimodal ISI distribution (**Fig.1F**) reflecting a propensity for bursting. By contrast, units were classified as putative INs if they had spike rates of ∼1-5Hz, short valley-to-peak and half width intervals, broad autocorrelations, and a predominantly unimodal ISI distribution (**Fig.1D-F**). All single units were required to have a refractory period ≥1ms. Units that had lower amplitude spikes and higher firing rates were classified as MUs (**Supplementary Fig.1A-B**). While this overall classification method is indirect and INs in particular have heterogeneous spiking properties (Tremblay et al., 2016), previous studies using human extracellular recordings have supported the classification of putative PYs and INs using similar metrics (Truccolo et al., 2011; Le Van Quyen et al., 2008).

### Sleep Spindle Detection and Analysis

Spindle detection was performed using a previously established method (Hagler et al., 2018) that is primarily based on the standard criterion of sustained power in the spindle band (**Fig.1G**). Each channel was bandpassed at 10-16Hz using an 8th order zero-phase frequency domain filter with transition bands equal to 30% of the cutoff frequencies. Absolute values of the bandpassed data were smoothed via convolution with a tapered 300ms Tukey window and median values of 10-16Hz band amplitudes were then subtracted from each channel to account for differences between channels. The data were then normalized by the median absolute deviation. Spindles were detected when the peaks in the normalized data exceeded 1 for at least 200ms, and onsets and offsets were marked when these amplitudes fell below 1. Putative spindles that coincided with large increases in lower (4-8Hz) or higher (18-25Hz) band power were rejected prior to analysis to exclude interictal epileptiform activity and broad spectrum artifacts, as well as 5-8Hz theta bursts, which may extend into the lower end of the spindle range (Gonzalez et al., 2018). Spindles from all channels of all patients were reviewed visually to confirm that epileptiform activity and artifacts were not present. Spindle frequency was calculated by dividing the number of zero crossings in the spindle band by two times the spindle duration. Coherence between co-occurring spindles was computed by finding the magnitude squared covariance of the 10-16Hz bandpassed spindle epoch. Spindle propagation velocity was calculated as 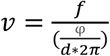, where *f* is the spindle frequency, *φ* is the phase offset between two waves, and d is the distance between the two waves.

### Unit Spike Timing Analysis

The percent of baseline spike rate during spindles for each unit detected on the same channel as the spindle was computed by multiplying 100 times the spike rate during all spindles on the unit’s channel divided by the spike rate of all epochs when no spindle was detected on the unit’s channel. Spindle phases of unit spikes were determined by computing the Hilbert transform of the 10-16Hz bandpassed signal and then finding the angle of the analytic signal at the times of the spikes. The circular mean angle was then computed for each unit (Berens, 2009).

### Paired Unit Spike Timing Analysis

Co-occurring spindles for each channel pair were identified if there was any spindle overlap. The first onset and last offset of the two spindles defined the co-occurring spindle epoch. Non-spindle epochs were the NREM periods when there was no spindle, with 100ms padded before the onset and after the offset of every spindle, detected on either channel for a given pair. Spiking of PY_1_-PY_2_, IN_1_-IN_2_, PY_1_-IN_2_, and IN_1_-PY_2_ during co-occurring spindles was compared to spiking during the non-spindle epochs as well as shuffled spikes during the co-occurring spindle intervals. Unit pair spiking was quantified by counting the number of spikes from one unit (e.g. PY_1_) during the 25ms preceding each spike from the second unit (e.g. PY_2_). Pairs with units detected on the same channel were excluded from analysis. Unit pairs with significantly increased co-firing within 25ms were tested for preference in the order of their firing. For each pair, the number of spikes during the 25ms window before vs. after the spikes of the other unit was compared. In addition, the correlation of co-firing within ±25ms between units was computed using the spike time tiling coefficient, which, unlike the correlation index, is independent of firing rates (Cutts and Eglen, 2014).

### Spatial Analysis of Spindles and Unit Spiking

The spatial layout of the recording array resulted in a highly variable number of contacts at different inter-contact distances. In order to approximately equalize the sample size in different distance bins, channel pairs were grouped progressively at increasing inter-contact distances until a minimum number was attained, and then a new bin was begun. For example, for determining the spatial fall-off of spindle co-occurrence, for each channel pair the number of co-occurring spindles with any overlap was determined, and then binned with at least 100 minimum channel pairs per distance bin. If there were subsequent pairs after 100 that had the same inter-contact distance then the values were included within that same bin. The distance values plotted show the average inter-contact distance for each bin. The same progressive binning method was used for the analysis of unit spike rates as a function of time and spindle coherence (minimum per bin=1000-5000), as well as spindle coherences (minimum per bin=100), spindle phase lags (minimum per bin=1000), unit spike rates (minimum per bin=50), unit co-firing significances (minimum per bin=50), and unit spike time tiling coefficients (minimum per bin=30) as a function of inter-contact distance.

### Analysis of Spindle Propagation

Visualization of sleep spindle propagation was done by z-scoring the 10-16Hz bandpassed data from each channel, which reduces effects of the average reference, as well as finding the phase of the 10-16Hz bandpassed data using the angle of the analytic signal. Prior to characterizing spatiotemporal patterns, epochs during which at least 20% of non-rejected channels were spindling were selected, and the rejected channels were spatially interpolated by performing a 2D biharmonic spline interpolation (MATLAB: *griddata)* of the analytic signal and then extracting the real signal. Spindle spatiotemporal patterns of propagation were characterized using the *NeuroPatt Toolbox* (Townsend and Gong, 2018), which uses optical flow estimation and singular value decomposition (SVD) to extract dominant spatiotemporal patterns from phase velocity vector field time series. This method is closely related to principal component analysis (PCA) as it reduces the dimensionality of the data to extract patterns that comprise the most variance. However, this method differs from PCA in that it extracts spatial modes that are vector fields, which represent spatiotemporal propagation patterns.

### Statistical Analyses

Differences in unit spike rates during spindles vs. baseline (non-spindle epochs during NREM) were evaluated using a one sample two-sided signed-rank test and differences between spike rates during spindles vs. baseline between unit types were tested using a two-sided rank-sum test. To test for a difference between the spindle phase distributions of PY and INs, a parametric Watson-Williams multi-sample test for equal circular means was used. To compare co-firing for each pair during co-occurring spindles vs. baseline, we randomly selected 1000 sets of non-spindle epochs, during which no spindles in either channel were detected, and which were matched in number and duration to the spindle co-occurrence epochs. The p-value was calculated as the percent of the 1000 sets of randomly selected non-spindle epochs that had more spikes from one unit in the 25ms preceding the spikes of the second unit. For example, for a given PY_1_-IN_2_ the number of PY_1_ unit spikes within 25ms preceding all IN_2_ spikes was counted. To compare co-firing for each pair during spindles vs. shuffled unit pair spiking during spindles, unit spikes during spindles were shuffled 1000 times and the p-value was calculated as described above. To test for spindle phase preferences of unit spiking, a Hodges-Ajne test was first used to determine if the distribution of the spindle phases of the spikes of each unit was non-uniform. Next, the spikes were randomly shuffled 1000 times and the Hodges-Ajne test was used to determine the 1000 p-values of the distribution of the spindle phases of the shuffled spikes. Finally, a unit was determined to have a significant phase preference if the p-value of the distribution was in the 5th percentile of the p-values of the shuffled distribution. The significance of ordered spiking of each unit pair was computed by comparing the proportion of the spikes from one unit occurring in the 25ms before vs. after the spikes from the other unit using a two-sided *χ*^2^ test of proportions (MATLAB: *chi2stat*) for all pairs with a minimum of 10 spikes during the ±25ms windows (Laurie, 2020). To compare spindle co-occurrence vs. chance, we compared the spindle co-occurrence density vs. the chance spindle co-occurrence density using a paired two-sided t-test. For each channel pair the chance spindle co-occurrence density was determined by randomly shuffling each channel’s spindles and inter-spindle intervals 100 times and finding the mean density of chance co-occurrences. When indicated, *α* levels were Bonferroni-corrected for multiple comparisons. All fits were approximated with a linear least squares regression, and for fits with R^2^<0.3, exponential least squares regressions were instead used if they met R^2^>0.3. If both fits met R^2^>0.3 then a linear fit was used unless the exponential fit had an R^2^ that was >25% larger. Fits are only shown for significant linear relationships or well-approximated exponential relationships. To test for the significance of a linear relationship, the significance of the correlation coefficient was used. To visualize the instantaneous phase of a propagating spindle, a cyclic color map was used (Thyng et al., 2016). To generate controls for the analysis of spatiotemporal propagation patterns, we shuffled the positions of the good channels prior to interpolation for each spindle. The Cohen’s d was calculated according to MATLAB: *computeCohen_d* (Bettinardi, 2020).

### Data and Code Availability

The data and custom code that support the findings of this study are available from the corresponding author upon reasonable request.

## Results

An average of 163 minutes (range: 120-200) of NREM sleep data was selected for analysis from recordings by the Utah Array implanted in supragranular layers II/III, and possibly as low as granular layer IV, of the superior or middle temporal gyrus in 4 patients (**Table 1**) with focal epilepsy undergoing monitoring for seizure localization prior to resection. 156 PYs, 39 INs, and 158 MUs were detected, classified, and analyzed. Therefore, among the single units, there were 80% PY and 20% IN. There were 496,269 spindles detected across 265 average referenced channels. The average and standard deviation spindle density per channel was 11.31±3.42 occurrences/minute, duration was 462.71±168.94ms, and oscillation frequency was 12.52±1.18Hz, which are consistent with previous intracranial studies in humans (Hagler et al., 2018; Mak-McCully et al., 2017). The average percent of spindles during which there was at least one spike on the same electrode from a PY was 8.94% (25th-75th percentiles: 1.76-14.35%) and an IN was 36.92% (20.85-52.76%).

### Spindles are associated with an increase in unit spike rates

Spike rates for each unit were quantified and analyzed during spindles detected on the unit’s channel and compared to a baseline period during which no spindles were detected on the unit’s channel. The median baseline spike rate of PYs was 0.11Hz (25th-75th percentiles: 0.01-0.24Hz) and INs was 0.74Hz (0.41-1.53Hz). The median spike rate during spindles for PYs was 0.24Hz (0.12-0.45Hz) and INs was 1.45Hz (0.68-2.55Hz). There was a significant increase in the median percent of baseline spike rate during spindles of 236% for PYs (25th-75th percentiles: 153-551%) and 183% for INs (127-264%, **Fig.1H**, Bonferroni-corrected p<0.0001, one-sample two-sided Wilcoxon signed-rank test). The increase in spike rate during spindles vs. baseline was similar for PY and IN among both patients with 1.0mm length probes and one patient with 1.5mm probes, while the other patient with 1.5mm probes had a greater increase in firing rate, perhaps because of a greater signal-to-noise ratio of spindles in superficial layer IV than deep layer III (Hagler et al., 2018).

### Unit spiking is phase-locked to the spindle with PY preceding IN

The circular mean spindle phase of spiking of PYs was 3.41rad (**Fig.1I**) and INs was 3.81rad (**Fig.1J**). There was a significant spindle phase preference of 21.29% of PYs and 56.41% of the INs (p<0.05, Hodges-Ajne test with bootstrapping significance). Since among the single units we detected, 80% were PY and 20% were IN, this suggests that 17% (100*0.2129*0.8) of neurons recruited with a significant phase preference were PYs and 11% (100*0.5641*0.2) were INs. For the units with significant spindle phase preferences, the circular mean spindle phase of spiking of PYs was 3.51rad (**Supplementary Fig.2A**) and INs was 3.94rad (**Supplementary Fig.2B**). There was a significant difference between the circular mean spindle phase angle distributions of PY spikes vs. IN spikes, with PY spiking preceding IN spiking (p<0.05, parametric Watson-Williams multi-sample test) for units with a significant phase preference. For spindles that range between 10-16Hz in frequency, this corresponds to a 4.02-6.44ms delay from PY to IN spiking, and for the average spindle frequency of 12.52Hz that we computed, this corresponds to a 5.47ms delay.

**Figure 2:**
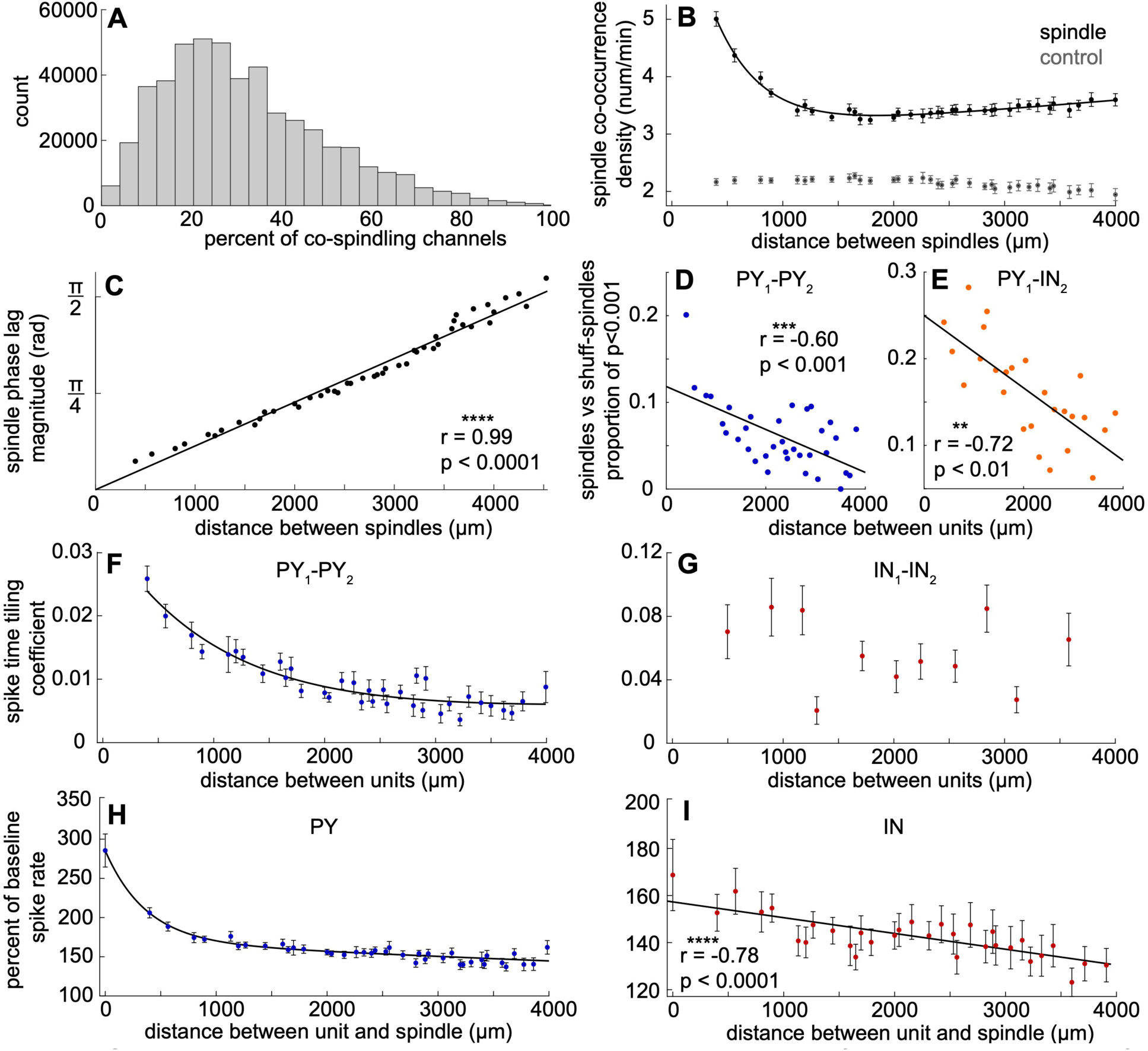
Spindling and spiking on a sub-centimeter scale. **A**, percent of co-spindling channels for all detected spindles. **B**, spindle co-occurrence density between channel pairs as a function of inter-contact distance. Black indicates measured spindle co-occurrence density and gray indicates chance spindle co-occurrence density if spindles occurred independently on different channels. **C**, magnitude of phase lag between co-occurring spindles as a function of inter-contact distance. **D-E**, mean proportion of unit pairs with significant co-firing within 25ms for spindles vs. shuff-spindles as a function of inter-contact distance for PY_1_-PY_2_ (**D**) and PY_1_-IN_2_ (**E**). **F-G**, spike time tiling coefficients (a measure for how much two spike trains are related) for PY_1_-PY_2_ (**F**) and IN_1_-IN_2_ (**G**) as a function of inter-contact distance. **H**, percent of baseline spike rate of PY for a co-located spindle (distance=0) or for spindles at progressively greater distances. **I**, same as **H** but for IN. Error bars show standard error of the mean. Fits for **B, F**, and **H** are two term exponential least squares regressions and fits for **C-E** and **I** are linear least squares regressions.

### Spindles preferentially co-occur within ∼1 mm

The spatial distribution of cortical spindles on a sub-centimeter scale has not been reported in humans. The median percentage of other channels spindling (given that at least one channel spindled) was 28.33% (25th-75th percentiles: 18.31-42.68%, **Fig.2A**). Spindle co-occurrence density (the frequency of spindle co-occurrence between two channels) was greatest at the smallest inter-contact distance of 400µm and decreased sharply until ∼1000µm, and then plateaued up to the maximum inter-contact distance of ∼4000µm (**Fig.2B**; R^2^=0.98, two term exponential least squares regression). For each channel pair the chance spindle co-occurrence density was computed by shuffling the spindles and inter-spindle intervals 100 times and then finding the mean co-occurrence density. The spindle co-occurrence density was greater than chance at all distances (Bonferroni-corrected p<0.0001, paired two-sided t-test, mean t=24.31, range=17.06-33.74, mean Cohen’s d=1.58, range=1.31-1.96).

### Sleep spindles propagate across the microarray with a characteristic velocity

There was a significant positive linear relationship between distance and the magnitude of the spindle phase lag between co-occurring spindles (**Fig.2C**, r=0.99, p<0.0001, significance of the correlation coefficient). Based on the magnitude of phase lag between co-occurring spindles as a function of distance, we estimated the spindle propagation velocity using the slope of the equation of the best fit linear regression, y=3.39e-4x+0.04, and the average spindle frequency of 12.52Hz. This calculation yielded a spindle propagation velocity of 0.23m/s.

### Unit pairs including PY co-fire preferentially at short distances

We next tested whether the proportion of unit pairs with significantly increased co-firing within 25ms during spindles vs. shuff-spindles was correlated with distance. There was a significant negative linear relationship for PY_1_-PY_2_ (**Fig.2D**, r=-0.60, Bonferroni-corrected p<0.001, significance of the correlation coefficient) and PY_1_-IN_2_ (**Fig.2E**, r=-0.72, Bonferroni-corrected p<0.01). To determine whether PY and IN have correlated firing that depends on distance, we evaluated PY_1_-PY_2_ and IN_1_-IN_2_ co-firing within ±25ms for all NREM periods using the spike time tiling coefficient, which is independent of firing rates (see **Methods**). PY_1_-PY_2_ correlations decreased with distance until ∼2000µm (**Fig.2F**, R^2^=0.88 two term exponential least squares regression). By contrast, IN_1_-IN_2_ correlations were larger overall and not dependent on distance at this scale (**Fig.2G**). The PY_1_-IN_2_ and IN_1_-PY_2_ correlations are not reported since the spike time tiling coefficient test cannot independently assess their correlations.

### Increased spindle-related spiking is maximal close to the detected spindle

In order to determine the spatial relationship between spindling and spiking, we evaluated the percent of baseline spike rate during spindles as a function of the distance between the unit channel and the spindle channel. PY unit spiking was greatest (∼285% of baseline) when the unit channel was spindling (i.e. distance of 0) and decreased with distance from the spindle channel until ∼1000µm (∼170% of baseline), at which point it decreased gradually (∼144% of baseline at 4000µm, **Fig.2H**, R^2^=0.94). IN unit spiking was also greatest when the unit’s channel was spindling (∼157% of baseline), and gradually decreased across space (∼130% of baseline at 4000µm) with a linear relationship (**Fig.2I**, r=-0.78, p<0.0001, significance of the correlation coefficient). Therefore, the gradual decrease from ∼1000-4000µm for PY was similar to the gradual decrease from ∼400-4000µm for IN, and at shorter distances from the spindle PY spiking was greatly increased.

### Spindles group unit pair co-firing within the window of STDP

We examined whether spindles organize unit pair spiking within the 25ms of STDP by computing peri-spike time histograms of spike counts, with the spikes of the first unit locked to t=0. We then plotted the spike rate of 1ms bins for 8026 PY_1_-PY_2_ pairs of units during spindles (**Fig.3A**) and non-spindle epochs during NREM sleep (**Fig.3B**), 580 IN_1_-IN_2_ during spindles (**Fig.3E**) and non-spindle epochs (**Fig.3F**), 2127 PY_1_-IN_2_ during spindles (**Fig.3G**) and non-spindle epochs (**Fig.3H**), and 2127 IN_1_-PY_2_ during spindles (**Fig.3K**) and non-spindle epochs (**Fig.3L**). For all types of unit pairs, the spindle distributions were shifted upward and there was a concentrated increase within ∼±25ms for spindle vs. non-spindle epochs, reflecting the overall and specific increases in spiking during spindles. To evaluate whether there was an increase in unit pair co-firing consistent with STDP, we compared the number of spikes from one unit that occurred within 25ms before the spikes of another unit for all possible pairs during spindles individually vs. 1000 shuffled non-spindle epochs matched in number and duration. There was a significant increase in unit pair spiking within 25ms for 25.54% of PY_1_-PY_2_ pairs, 60.69% of IN_1_-IN_2_, 50.16% of PY_1_-IN_2_, and 49.32% of IN_1_-PY_2_, during spindles vs. non-spindle epochs (p<0.001, bootstrapped significance, **Table 2A**).

**Figure 3:**
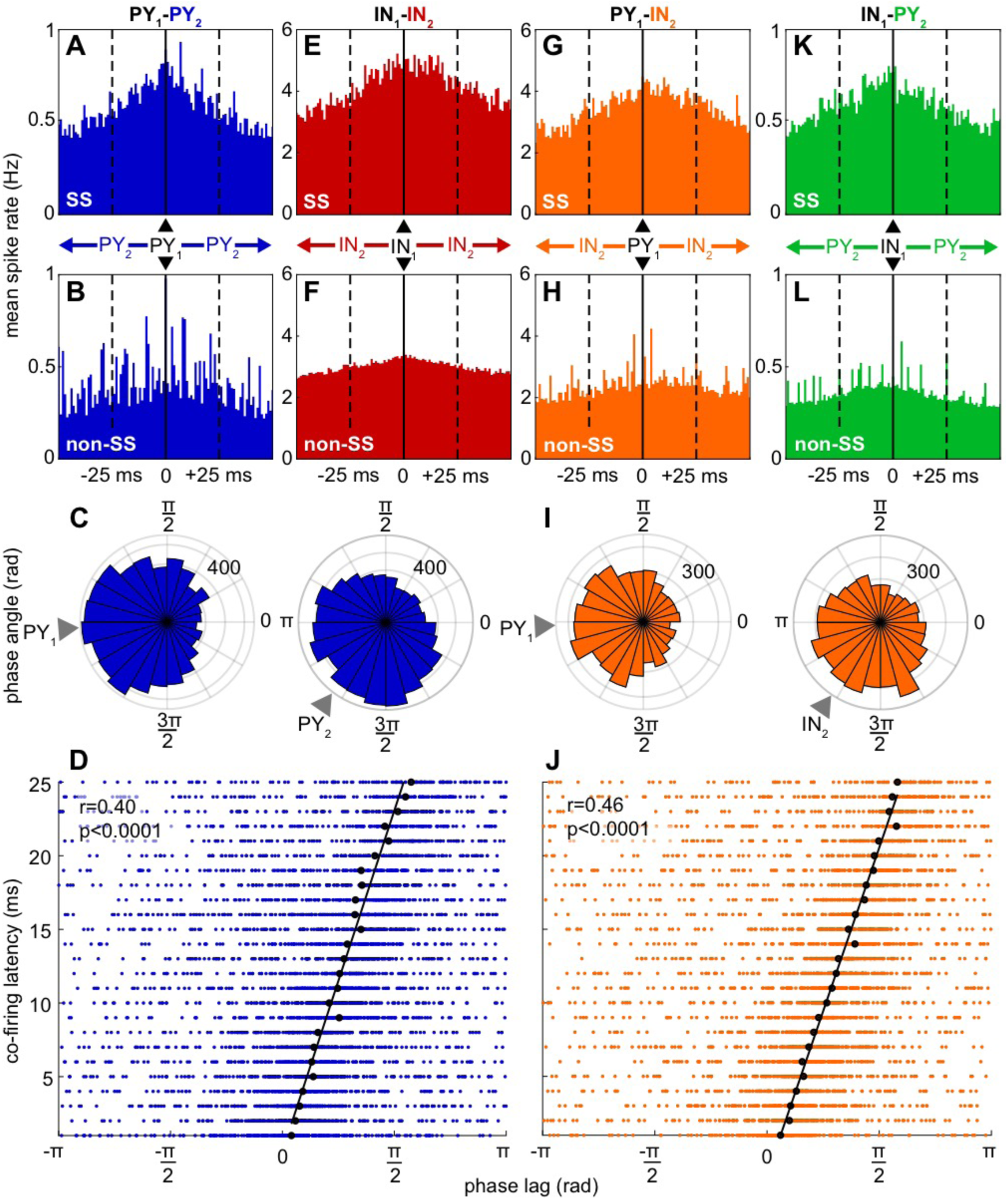
Unit pair co-firing during spindles. **A**, in all pairs of PY recorded from different electrodes, the firing of one (PY_2_) is plotted relative to the time of the other (PY_1_, at t=0) during spindles (**A**) and between spindles (**B**). Data are averaged across 1ms bins. Solid vertical line shows t=0. Dashed vertical lines show the ±25ms interval where paired pre- and post-synaptic spiking facilitates STDP. **C**, polar histograms showing co-located spindle phases of PY_1_ and PY_2_ spikes. The two units of each pair and their co-located spindles were detected on separate channels in all cases. Gray triangles indicate circular means. **D**, co-firing latency is significantly correlated with the phase lag between the local spindles co-located with the co-firing units. Black circles indicate circular means for each 1ms binned latency and black line indicates circular-linear best fit of these means. **E-F**, same as **A-B**, except with IN_1_-IN_2_. **G-J**, same as **A-D** except with PY_1_-IN_2_. **K-L**, same as **A-B** except with IN_1_-PY_2_.

**Table 2:** Significance of paired and ordered unit co-firing. **A-C**, percent of unit pairs with significantly increased (p<0.001) co-firing between unit pairs within 0-25ms during spindles vs. non-spindles (**A**), spindles vs. shuff-spindles (**B**), and both (**C**). **D**, percent of unit pairs in **C** with significant order preference of co-firing within 25ms (p<0.05, two-sided *χ*^2^ test of proportions). Shuff-spindles=sleep spindles with shuffled spikes.

| Unit Pair (1→2) | A. Paired unit co-firing: spindles vs. non-spindles | B. Paired unit co-firing: spindles vs. shuff-spindles | C. Paired unit co-firing: Both | D. Ordered spiking |
| --- | --- | --- | --- | --- |
| PY<sub>1</sub>-PY<sub>2</sub> | 25.54% | 7.03% | 6.83% | 26.60% |
| IN<sub>1</sub>-IN<sub>2</sub> | 60.69% | 37.93% | 34.66% | 18.00% |
| PY<sub>1</sub>-IN<sub>2</sub> | 50.16% | 16.93% | 15.94% | 23.14% |
| IN<sub>1</sub>-PY<sub>2</sub> | 49.32% | 14.01% | 13.12% | 28.78% |

### Unit pair co-firing within the window of STDP during spindles is not solely due to increased spike rates

Increased co-firing within the 25ms could simply be due to the increase in spike rates during spindles (**Fig.1H**). However, unit co-firing distributions have steeper slopes within 25ms (**Fig.3**). To test if unit pair co-firing distinctly increases within 25ms, we compared paired unit spiking within 25ms during spindles vs. the same spindles with shuffled spikes (shuff-spindles). There was a significant increase in paired spiking during spindles vs. shuff-spindles for 7.03% of PY_1_-PY_2_, 37.93% of IN_1_-IN_2_, 16.93% of PY_1_-IN_2_, and 14.01% of IN_1_-PY_2_ (p<0.001, bootstrapped significance, **Table 2B**). About 95% of these unit pairs also increased co-firing significantly during spindles vs. non-spindles (6.83% of PY_1_-PY_2_, 34.66% of IN_1_-IN_2_, 15.94% of PY_1_-IN_2_, and 13.12%, p<0.001, bootstrapped significance, **Table 2C**). Therefore, the increase in unit pair co-firing during spindles is due not only to the overall rate increase, but also to a specific grouping within 25ms, presumably associated with the phase-clustered firing. Overall, about a quarter of PY pairs and half of IN pairs increase co-firing during spindles, and of these about a quarter of PY pairs and three-quarters of IN pairs further increased co-firing beyond that expected from the general increase in firing rate during spindles.

### Unit pairs show ordered co-firing during spindles

In the canonical model, STDP is an order-dependent process that can lead to LTP or LTD (Feldman, 2012). Therefore, we investigated whether unit pairs with significantly increased co-firing within 25ms for both spindles vs. non-spindles and spindles vs. shuff-spindles had a preferred order of firing within this window. Of 282 PY_1_-PY_2,_ 200 IN_1_-IN_2,_ 242 PY_1_-IN_2,_ and 278 IN_1_-PY_2_ pairs with ≥10 co-firing spikes, 26.60% of PY_1_-PY_2_, 18.00% of IN_1_-IN_2_, 23.14% of PY_1_-IN_2_, and 28.78% of IN_1_-PY_2_ had a preferred order of spiking (two-sided *χ*^2^ test of proportions, p<0.05, **Table 2D**).

### Co-firing lags across electrodes are strongly correlated with co-located spindle phase-lags

Since unit spiking is locked to spindle phase and there is an increase in co-firing during spindles, we tested whether co-firing was also locked to spindle phase. For PY_1_-PY_2_, when PY_2_ fired within 25ms following PY_1_, PY_1_ spikes had a circular mean phase of 3.2rad and PY_2_ spikes had a circular mean phase of 4.1rad (**Fig.3C**). Likewise for PY_1_-IN_2,_ PY_1_ had a circular mean phase of 3.2rad and IN_2_ had a circular mean phase of 4.1rad (**Fig.3I)**. There was no significant difference between spindle phase of PY_1_ in PY_1_-PY_2_ vs. PY_1_ in PY_1_-IN_2,_ or between spindle phase of PY_2_ in PY_1_-PY_2_ vs. IN_2_ in PY_1_-IN_2_ (p>0.05, parametric Watson Williams multi-sample test). When we analyzed co-firing latency vs. spindle phase lag there was a significant circular-linear relationship for both PY_1_-PY_2_ (**Fig.3D**, r=0.40, p<0.0001, significance of the circular-linear correlation coefficient) and PY_1_-IN_2_ (**Fig.3J**, r=0.46, p<0.0001). In sum, when two cells recorded by different electrodes fire within 25ms of each other during spindles, the latency between their spikes is highly correlated with the phase lag between the spindles recorded by the two electrodes.

### Co-firing across electrodes is strongly correlated with co-located spindle coherence

We next tested if spindle coherence between electrodes within the 10-16Hz band was associated with co-firing recorded by those electrodes (schematic in **Fig.4A**). The magnitude squared coherence of local LFP within the 10-16Hz band during spindles was greater at all distances compared to randomly selected NREM epochs matched in number and duration (**Fig.4B**; Bonferroni-corrected p<0.0001, two-sample two-sided t-test, mean t=110.36, range=31.41-207.63, mean Cohen’s d=0.54, range=0.38-0.71). High levels of co-firing between units recorded by different electrodes was restricted to high levels of spindle coherence (>∼0.95) between those electrodes (**Fig.4C-J)**. This co-firing was mainly at short lags (<∼10ms), and was observed for all pair types. For example, the mean co-firing rate at lags <10ms by PY-PYs increased by ∼50% when coherence was very high (>0.95), versus when the coherence was lower (<0.90), and by ∼200% compared to baseline NREM periods between spindles (**Fig.4D**; Bonferroni-corrected p<0.0001, two-sample two sided t-test, t=10.45 and 38.29, respectively, Cohen’s d=1.10 and 3.77, respectively). For IN-INs, the corresponding increases were ∼50% and ∼120% (**Fig.4F**; Bonferroni-corrected p<0.0001, two-sample two-sided t-test, t=13.64 and 36.06, respectively, Cohen’s d=1.30 and 3.56, respectively; see **Supplementary Table 1** for additional details). Thus, short latency unit co-firing depends critically on spindle coherence being close to 1.

**Figure 4:**
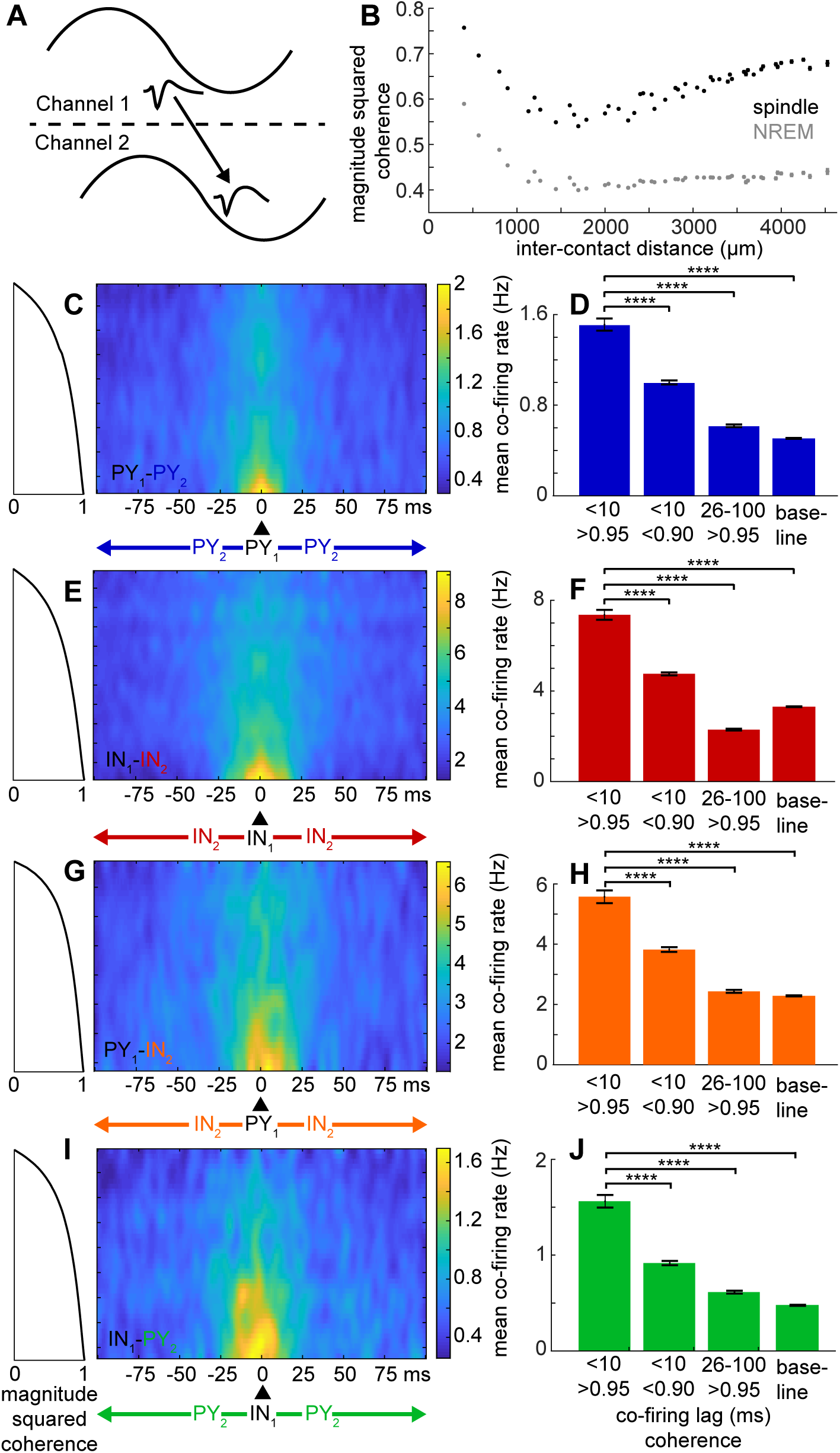
Spindle coherence and unit pair co-firing. **A**, schematic portraying one cycle of co-occurring spindles and co-firing between units on two separate channels. **B**, effect of distance on magnitude squared coherence in the 10-16Hz band during co-occurring spindles (black) and during randomly selected NREM epochs matched in number and duration (gray). **C**, PY_1_-PY_2_ co-firing rates as a function of spike lags and spindle coherence. **D** mean co-firing rates in **C** for shorter spike lags (<10ms) and higher coherence (>0.95) in the left bar, compared to shorter lags and lower coherence (<0.90), longer lags (26-100ms) and higher coherence, and shorter lags during baseline NREM periods in between spindles. **E-J**, same as **C-D**, except for IN_1_-IN_2_ (**E-F**), PY_1_-IN_2_ (**G-H**), and IN_1_-PY_2_ (**I-J**). Co-firing heatmaps were smoothed with a 2D Gaussian filter with *α*=2. Error bars show SEM. Bonferroni-corrected ****p<0.0001.

### Spindles propagate across the microarray in multiple patterns

Different waves within an individual spindle could exhibit multiple patterns of propagation (**Fig.5A-F; Video 1**). For example, one spindle wave had a circular propagation pattern, based on its z-score normalized amplitude (**Fig.5A,E**) and phase (**Fig.5B,F**), and in subsequent wave cycles showed a planar propagation pattern (**Fig.5C-F**). We used the MATLAB: *NeuroPatt Toolbox* (see **Methods**) to find spatiotemporal modes, represented as phase velocity vector fields, that explained the greatest percent variance of the phase velocity vector time series for each spindle (**Fig.6**). Controls were generated by shuffling the positions of the good channels prior to interpolation and spatiotemporal analysis for each spindle. The percent explained variances of modes 1 and 2, i.e. those with the greatest percent explained variance, were greater for spindles vs. shuffled controls in all subjects (**Fig.6A-B**, p<0.0001, two-sided signed-rank test), providing confirmation of propagating spindles. There were a variety of propagation patterns within and across patients and spindles (representative examples of mode 1 in **Fig.6C**), demonstrating that spindles have multiple patterns of propagation on a 4mm scale.

**Figure 5.**
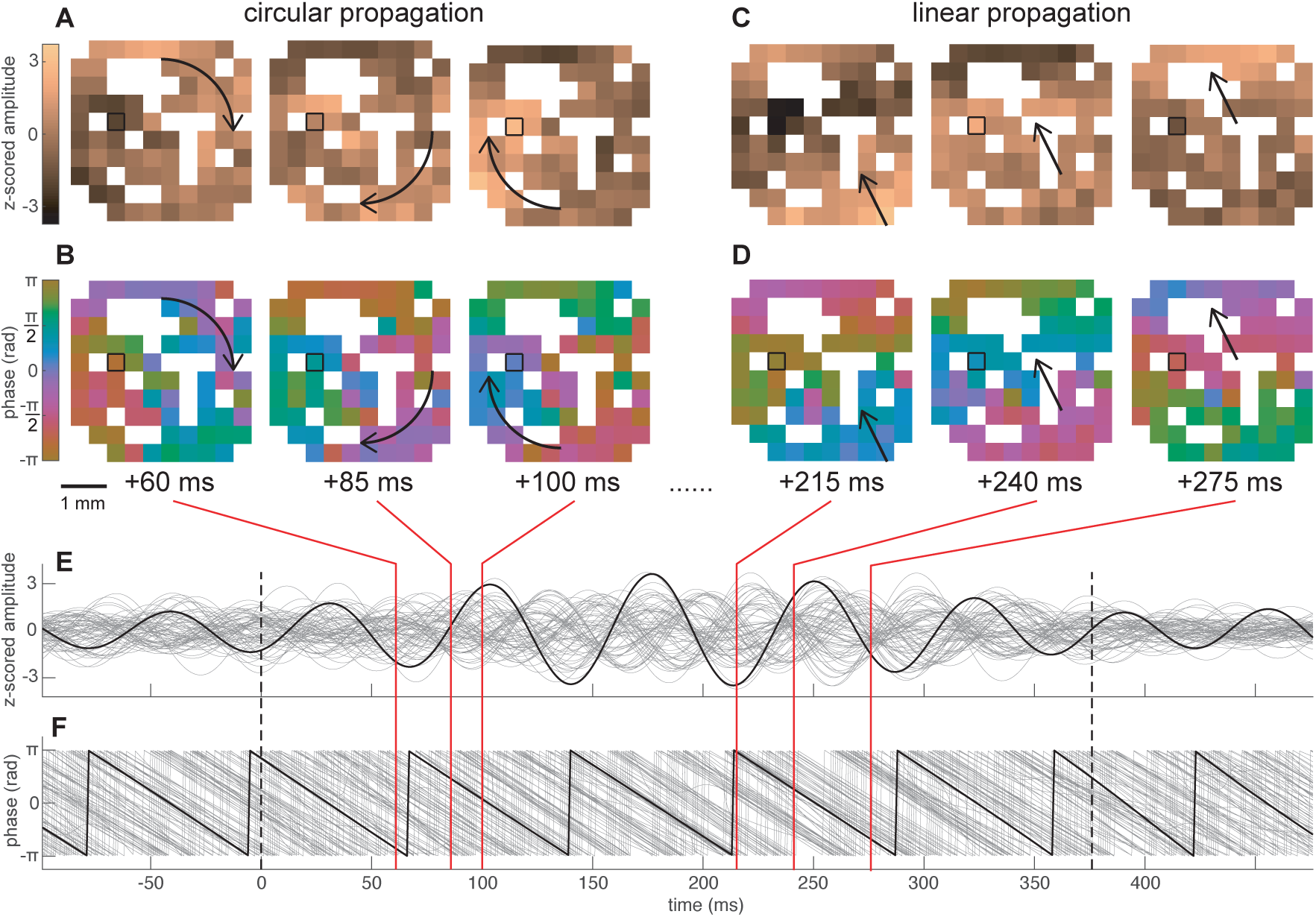
Sleep spindles propagate on a sub-centimeter scale. **A-B**, circular propagation of a spindle wave depicted in 3 frames of the instantaneous z-score normalized amplitude (**A**) and instantaneous phase (**B**). A cyclic colormap was used to show phase. **C-D**, same as **A-B** but for planar propagation of a different wave within the same spindle. White spaces indicate non-existent or bad channels. The channel outlined in black corresponds to the channel on which the spindle was detected. Arrows indicate approximate trajectory of propagation. **E**-**F** traces of z-score normalized amplitude (**E**) and phase (**F**) of the same spindle in **A-D**. The black trace corresponds to the channel on which the spindle was detected and gray traces show the rest of the channels. Red lines extending from **E** and **F** show the times of the frames in **A-D**. See corresponding **Video 1**.

**Figure 6.**
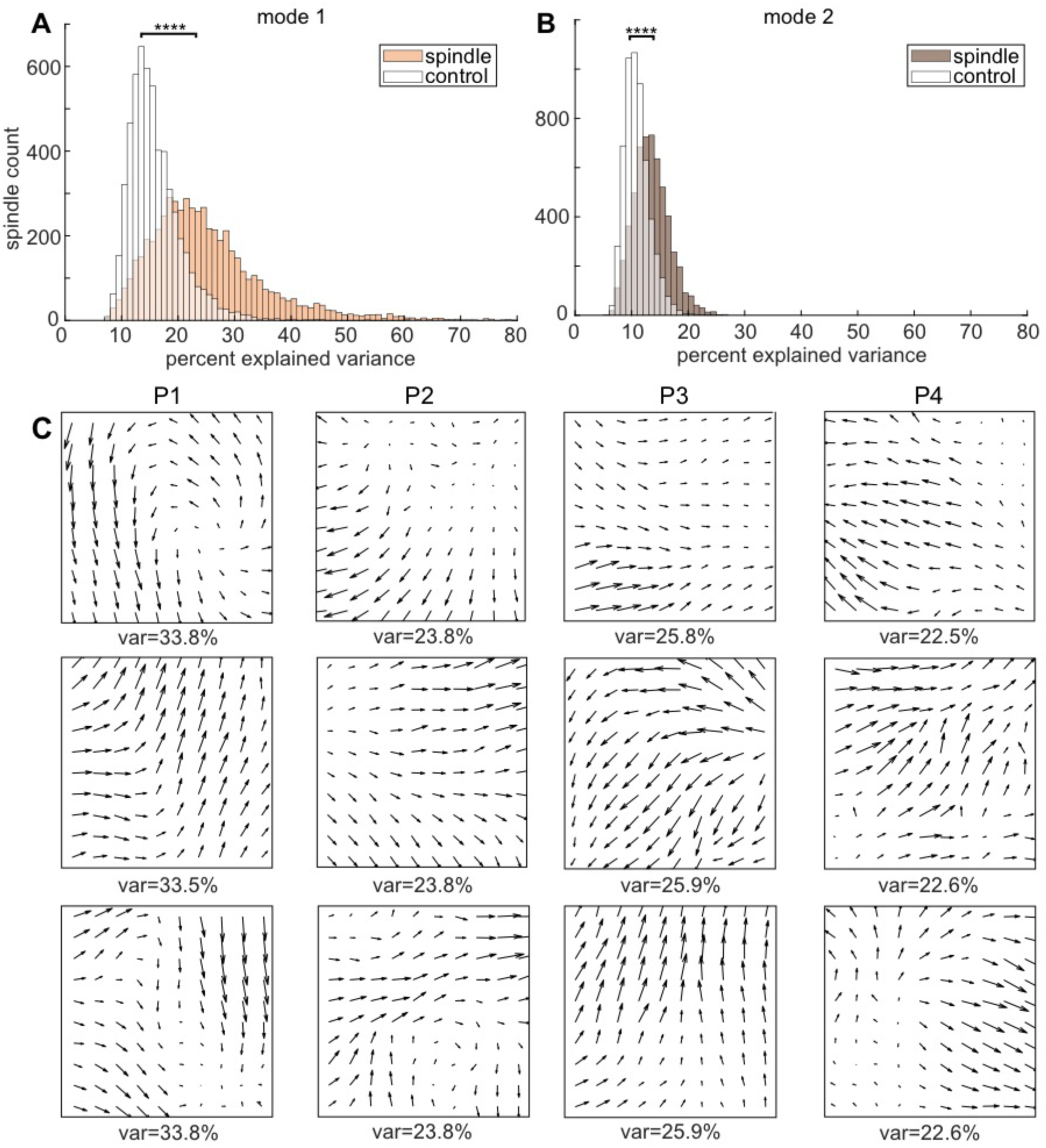
Spatiotemporal propagation patterns of spindles. **A**, percent explained variance (var) for most dominant spatiotemporal mode (mode 1) across spindles (tan) and shuffled controls (white). **B**, same as **A** except for mode 2. **C**, three representative examples of mode 1 phase velocity vector fields showing individual spindle propagation patterns for each patient (P1-4). ****p<0.0001.

### Distinct spindle propagation patterns are associated with distinct unit firing patterns

The above spindle propagation analysis analyzed each spindle independently, and so the patterns could also change across spindles. In order to analyze co-firing it was necessary to identify consistent propagation patterns. Thus, a second analysis was performed on a concatenation of all spindles from each patient. Again, distinct modes were consistently found as in **Fig.6C**. PY_1_-PY_2_ co-firing was then quantified independently for modes 1 and 2 of the concatenated spindles for each patient when SVD component scores exceeded the 75^th^ percentile, representing periods when each mode was most prominent. Among 205 pairs where PY_2_ co-fired within 25ms following PY_1_ at least 10 times between modes 1 and 2, 183 pairs (89.27%) had a significant difference in co-firing for modes 1 vs. 2 (two-sided *χ*^2^ test of proportions, *α*=0.05). Thus, distinct spindle propagation patterns are associated with different sequences of unit co-firing.

## Discussion

We identified the spatiotemporally-patterned inter-relations of LFPs and neuronal firing during human sleep spindles over a 10×10 array of microelectrodes at 400µm pitch. Firing of individual putative cortical PYs and INs increased during spindles, with an additional increase at certain spindle phases. Co-firing within 25ms of cell pairs recorded on different microelectrodes, a pre-requisite for STDP, also increased during spindles in a phase-locked manner. Co-firing of cells and co-occurrence of spindles were greatest at interelectrode separations <1mm but extended over the entire array. The average lag between co-firing cells and phase-lag between coherent spindles increased linearly with distance. Conduction speed and the spindle phase precedent of PY cells over IN were consistent with direct cortico-cortical spindle propagation. Co-firing at the shortest latencies was concentrated to electrode pairs with highly coherent spindles. Multiple two-dimensional spindle propagation patterns and associated distinct co-firing patterns occurred across each array, intermixed within and between spindles. Overall, these microphysiological mechanisms may support and organize memory consolidation by creating the necessary conditions for STDP and activating spatiotemporal networks through travelling spindles.

In contrast to Andrillon et al. (2011) who found no increase in firing during spindles in humans, we found a doubling. This difference could be because they sampled medial limbic cortex whereas we sampled lateral temporal cortex. Further, they correlated unit spiking detected by microwires with LFP recorded by a macroelectrode contact ∼4mm away, whereas we correlated unit spiking with LFP recorded from the same contacts, and/or because they recorded from all cortical layers whereas we only recorded from supragranular and possibly granular layers. Indeed, the increase in unit firing during spindles was ∼50% smaller at a distance of ∼4mm, and it has been previously shown that spindle-phase modulation of high gamma in humans and of unit firing in rodents is greater in supragranular vs. infragranular layers (Hagler et al., 2018; Peyrache et al., 2011). Thus, our recordings focused on the most responsive layers. We furthermore found that PY and IN spiking was locked to the phase of individual spindle waves, which is consistent with what has been previously shown for PY and IN in rodents (Peyrache et al., 2011) and for units without cell-type classification in humans (Andrillon et al., 2011), and suggests a mechanism whereby spindles specifically coordinate co-firing beyond a mere general tonic increase in firing.

Our study provides the first direct evidence that the essential pre-condition for STDP, unit pair co-firing within 25ms, is met in the human cortex during spindles in NREM sleep. A major issue in understanding how SDTP could occur under normal conditions is the large number of repetitions, up to hundreds, that are needed to produce long-term changes (Wittenberg and Wang, 2006). In a normal night’s sleep, ∼1000 spindles will occur at most cortical sites, and each spindle has ∼10 cycles, so there are ample opportunities for STDP repetitions. Furthermore, spindle frequency lies within the repetition rate for which such pairings are effective (Feldman, 2012). In addition, memory-related cortical input from the hippocampus associated with ripples may be available on multiple spindle peaks often seen in the posterior hippocampus, phase-locked with cortical spindles (Jiang et al., 2019a; Staresina et al., 2015).

In canonical PY_1_-PY_2_ STDP, pre-before-postsynaptic spiking leads to LTP and the reverse leads to LTD (Feldman, 2012). Plasticity underlying memory consolidation may involve both (Clopath, 2012). Increased co-firing within 25ms for PY_1_-PY_2_, PY_1_-IN_2_, IN_1_-PY_2_, and IN_1_-IN_2_ was found during co-occurring spindles up to >4mm, consistent with the proposition that co-occurring spindles may provide a context for transcortical plasticity (Muller et al., 2016). Many of the pairs with increased co-firing had a preferred order of spiking, which could support unidirectional plasticity. Human slice recordings have shown that local excitatory connectivity between layers II/III PYs is 13-18% (Peng et al., 2019), substantially higher than in mice (Seeman, 2018). However, most co-firing within 25ms is probably by unit pairs that are not directly connected but belong to the same local network. In some cases this is directional, however there could be multiple pathways between co-firing units, some in one direction and others in the opposite direction, and network tuning would involve strengthening some routes and weakening others.

Human magnetoencephalography (Dehghani et al., 2011) and intracranial macroelectrode (Piantoni et al., 2017; Nir et al., 2011) recordings have shown that spindles, once thought to be a global phenomenon, are often focal on a centimeter scale. Human laminar recordings have furthermore shown that spindles localize to specific cortical layers (Hagler et al., 2018; Halgren et al., 2018); however the lateral extent of spindles in human cortex has not been reported on a sub-centimeter scale. We show that spindle co-occurrence and coherence in human cortex peaks at the shortest inter-contact distance of 400µm, decreasing sharply to a plateau at ∼1000µm. Since an average reference would eliminate spindles that are equal across all leads, there may also be co-occurrence at a larger scale, and indeed, asymptotic co-occurrence and coherence exceeded chance. Taken together, the data indicate that the cortical extent engaged by spindles can range from a few columns to much of the cortex.

Spindles propagated within the microgrid at ∼0.23m/s. This is within the range of or slightly lower than previously reported intracortical axon conduction velocities, including for layers II/III, of 0.28m/s and 0.15-0.44m/s in rat visual cortex (Lohmann and Rörig, 1994; Murakoshi et al., 1993), 0.35-0.45m/s in rat neocortex (Telfeian and Connors, 2003), and 0.35m/s in cat visual cortex (Hirsch and Gilbert, 1991). The true axonal conduction velocity of spindles may be faster than 0.23m/s because our calculation assumes a direct path of travel and does not take synaptic delays into account. This velocity is much slower than what was reported by Muller et al. (2016) for human cortical spindles (3-9m/s) and Halgren et al. (2019) for human cortical alpha (0.91m/s), both using ECoG recordings, presumably because they were measuring fast conduction via myelinated fibers passing through the white matter, whereas we were measuring slow conduction via unmyelinated fibers within the cortical gray matter.

Direct cortico-cortical propagation of spindles is at odds with the common conception of cortical spindles being driven from the thalamus. In cats, the thalamus continues to spindle after cortical removal, but the cortex does not spindle after disconnection from the thalamus (Contreras et al., 1997). In mice, rhythmic optogenetic activation of the thalamic reticular nucleus triggers spindles (Halassa et al., 2011). In humans, thalamic spindles occur more frequently and begin before cortical, and in rare cases show tight phase locking with thalamus leading the cortex (Mak-McCully et al., 2017). Thus, spindles are thought to originate thalamically and project cortically (De Gennaro and Ferrara, 2003). Thalamocortical projections in mice to both primary sensory and limbic cortices drive INs more strongly and at shorter latencies than PYs (Cruikshank et al., 2007; Delevich et al., 2015). Thus, our finding that PY spiking *precedes* IN spiking is not consistent with cortical spindles in humans being mainly driven by the thalamus. Rather, it is possible that while cortical spindles in humans are initially driven by the thalamus, intrinsic cortical circuits may subsequently amplify and spread the spindle. Local generation seems plausible because the thalamic mechanism underlying spindle generation involves reciprocal connections between excitatory and inhibitory cells, and activations of h and T currents (Destexhe and Sejnowski 2003; McCormick et al., 2015), all of which are present in human supragranular cortex (Kalmbach et al., 2018). Furthermore, the consistently higher spindle frequency in the human thalamus compared to cortex is hard to explain if cortical spindles are all directly driven by the thalamus (Mak-McCully et al., 2017). This thalamocortical frequency difference increases over the course of a spindle, as the spindle spreads across the cortex, and is correlated with the amount of such spread (Gonzalez et al., unpublished). Direct cortico-cortical spindle propagation may be necessary in humans, who have ∼1400 cortical neurons for every thalamocortical cell (calculations based on cell counts in Azevdo et al., 2009; Xuereb et al., 1991).

In summary, we show here that human cortical neurons have a strong increase in firing during spindles, both tonically and at particular spindle phases. This is associated with greatly enhanced co-firing by cells recorded by different microelectrodes separated by ∼0.4-4mm at delays <25ms, a precondition for STDP. Within this microdomain, spindles propagate in multiple consistent patterns, achieving high coherence between contacts which is strongly associated with short-latency co-firing by the units they record. Multiple patterns of wave propagation occur both within and between spindles, and are each associated with distinct sequences of co-firing. Since the co-firing latency is highly correlated with the phase lag between spindles, these data together suggest that travelling spindles may organize spatiotemporal sequences of neuronal firing to modulate their synaptic strengths via STDP.

## Acknowledgements

Supported by NIMH (RF1 MH117155, T32 MH020002), ONR-MURI (N00014-16-1-2829), NIBIB (R01 EB009282), and the Kavli Institute for Brain and Mind. We thank Adam Niese, Burke Rosen, Christopher Gonzalez, Daniel Cleary, Erik Kaestner, Jacob Garrett, Sophie Kafjez, Xi Jiang, Yihan Zi, and Zarek Siegel for their support.

