## Supplementary Information for "Travelling spindles create necessary conditions for spike-timing-dependent plasticity in humans"

*Multi-unit spiking during spindles.*

The average percent of spindles during which there was at least one spike on the same electrode from a MU was 52.18% (26.62-81.82%). The median baseline spike rate of MUs was 1.41Hz (25th-75th percentiles: 0.43-3.60Hz). The median spike rate of MUs during spindles was 2.67Hz (0.98-5.98Hz). There was a significant increase in the median percent of baseline spike rate during spindles of MUs of 164% (134-223%, p<0.0001, one-sample Wilcoxon signed-rank test; **Supplementary Fig.1C**). The circular mean spindle phase of spiking of MUs was 3.41rad (**Supplementary Fig.1D**). There was a significant spindle phase preference for of MUs (p<0.05, Hodges-Ajne test with bootstrapping significance). There was a significant spindle phase preference 49.37% of the MUs (p<0.05, Hodges-Ajne test with bootstrapping significance). For the MUs with significant spindle phase preferences, the circular mean spindle phase of spiking was 3.38rad (**Supplementary Fig.2C**). We examined whether spindles organize MU-MU spiking within the 25ms of STDP by computing peri-spike time histograms of spike counts, with the spikes of the first unit locked to t=0. Based on analysis of 8930 pairs, MU-MU co-firing was increased within 25ms (**Supplementary Fig.3**).

*Unit pair co-firing, except for $PY_1$-$PY_2$, is greatest when there are co-localized spindles.*

We tested for significant co-firing of unit pairs for epochs during which one unit was detected on a channel participating in spindles and the other unit was detected on a channel not participating in spindles vs. the shuffled spikes of both units for the corresponding conditions (i.e. spindles+non-spindles vs. shuff-spindles+shuff-non-spindles). There was a similar percentage of significant $PY_1$-$PY_2$ compared to spindles vs. shuff-spindles (7.33% vs. 7.03%), but the percentages of significant $IN_1$-$IN_2$, $PY_1$-$IN_2$, and $IN_1$-$PY_2$ were much less (7.41% vs. 37.93%, 3.76% vs. 16.93%, 1.50% vs. 14.01%, respectively, **Supplementary Table 2**). This shows that the propensity of unit pairs containing at least one IN to co-fire tends to require that spindles co-occur at both locations.

| | **<10 ms, >0.95 coh** | **<10 ms, <0.90 coh** | **26-100 ms, >0.95** | **baseline** |
|---|---|---|---|---|
| **$PY_1$-$PY_2$** | 1.51±0.55 Hz | 1.00±0.45 Hz | 0.62 ±0.31 Hz | 0.51±0.49 Hz |
| **$IN_1$-$IN_2$** | 7.36±2.46 Hz | 4.76±1.92 | 2.29±1.26 Hz | 3.29±0.93 Hz |
| **$PY_1$-$IN_2$** | 5.57±2.15 Hz | 3.82±1.81 Hz | 2.44 ± 1.33 Hz | 2.27±1.04 Hz |
| **$IN_1$-$PY_2$** | 1.56±0.99 Hz | 0.92±0.79 Hz | 0.61±0.60 Hz | 0.50±0.42 Hz |

Supplementary Table 1: Unit pair co-firing rates according to spindle phase lag and coherence. Mean and standard deviation unit pair co-firing rates during co-occurring spindles for shorter co-firing lags (<10ms) and higher spindle coherence (>0.95), shorter lags and lower coherence (<0.90), longer lags (26-100ms) and higher coherence, and shorter lags during baseline NREM periods in between spindles.

| Paired unit co-firing: spindles+non-spindles vs. shuff-spindles+shuff-non-spindles |
|---|
| 7.33% |
| 7.41% |
| 3.76% |
| 1.50% |

<u>Supplementary Table 2: Significance of paired unit co-firing when only one channel is spindling.</u> Percent of unit pairs with significantly increased ($p<0.001$) co-firing within 0-25ms for real vs. shuffled spikes of unit pairs during times when one unit's channel is spindling and the other is not.

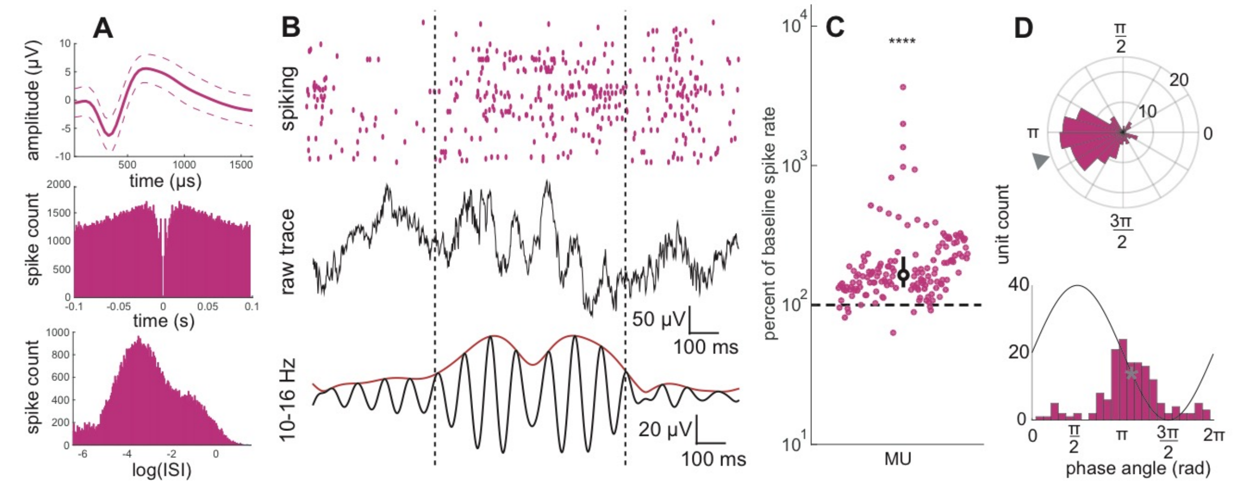


<u>Supplementary Figure 1: Multi-unit classification, spiking during spindles, and spindle phase preference.</u> **A**, average and standard deviation MU spike waveform (top), spike autocorrelation (middle), and ISI distribution (bottom). **B**, raw and 10-16Hz bandpassed traces of a sleep spindle with raster plot of associated MU spiking. **C**, MU spike rates during spindles as log-percent of baseline firing. The spindle firing rate of each unit was computed as the firing rate across all spindles. The baseline firing rate was computed as the firing rate during epochs without spindles. Each colored circle shows the mean firing rate of one unit. White circle shows the median value and error bars show the 25th and 75th percentiles. Dashed horizontal line shows 100% percent, which is equal to the baseline non-spindle spike rate. **D**, polar and non-polar histograms show circular mean spindle phases of MU spiking. One cycle of a spindle is superimposed on the non-polar histogram to visualize the phase-spike timing relationship. Gray triangle on polar histograms and gray asterisk on non-polar histogram shows circular mean. ****$p<0.0001$. ISI=inter-spike interval, MU=multi-unit.

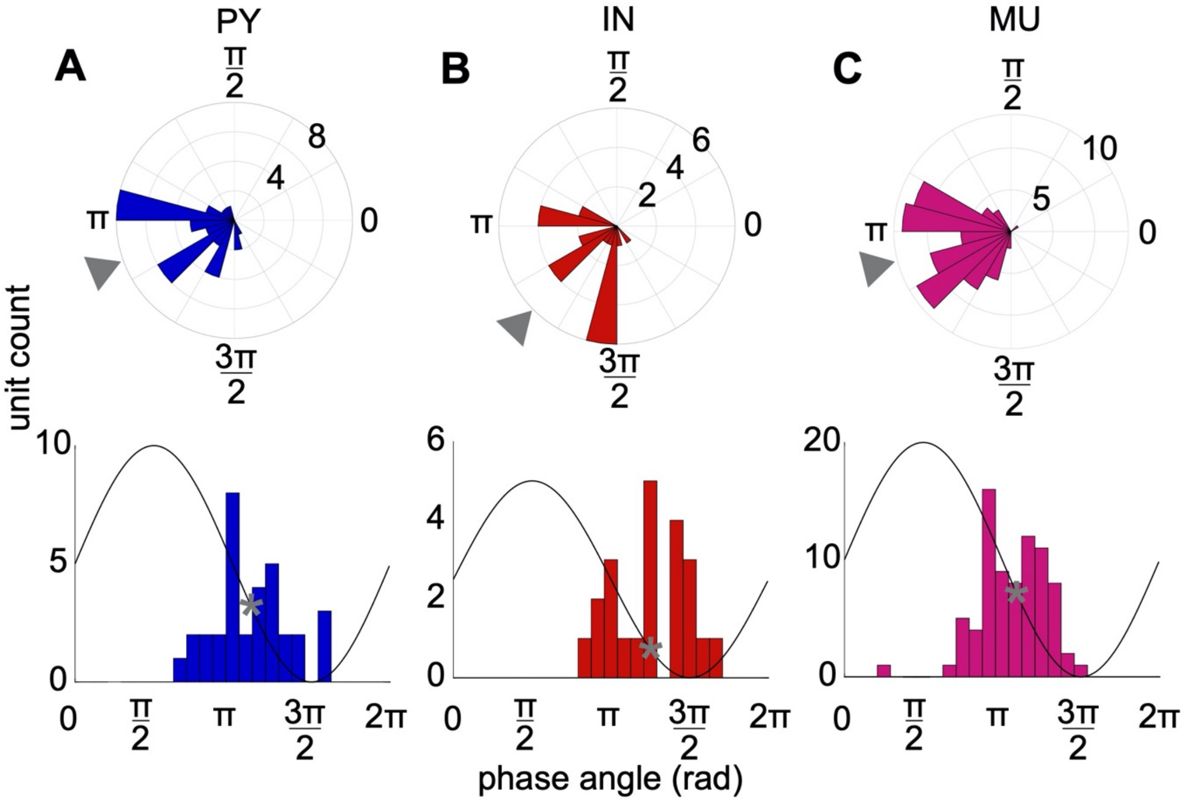


<u>Supplementary Figure 2: Spindle phase preference of units with statistically significant non-uniform spiking distributions.</u> **A,** polar and non-polar histograms show circular mean spindle phases of spiking of PYs (**A**), INs (**B**), and MUs (**C**) with statistically significant non-uniform spike distributions (see **Methods**). One cycle of a spindle is superimposed on non-polar histograms to visualize the phase-spike timing relationship. Gray triangles on polar histograms and gray asterisks on non-polar histograms show circular means.

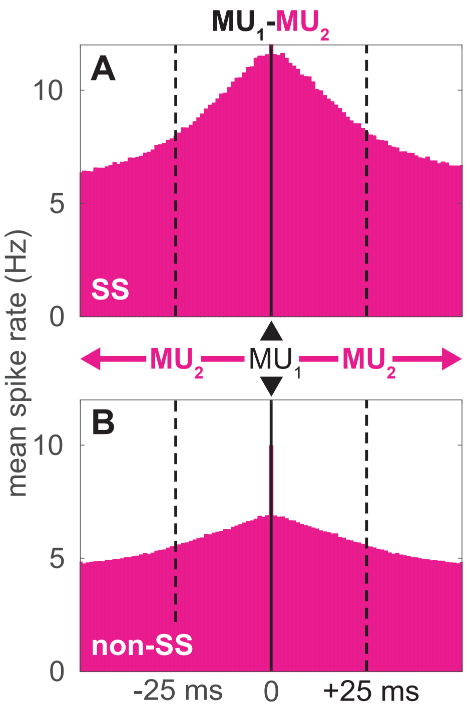


Supplementary Figure 3: Paired MU spiking during spindles and non-spindle epochs. **A-B**, peri-spike time histogram of multi-unit spikes locked at t=0 to each spike from a paired MU for all MU-MUs during spindles (**A**) and non-spindle epochs (**B**). Solid vertical line shows t=0. Dashed vertical lines show the ±25ms interval where pre-synaptic spiking followed by post-synaptic spiking facilitates STDP.
